# α -tACS Modulates Reward-Dependent Pupil Responses and Corticostriatal Connectivity

**DOI:** 10.64898/2026.01.30.702963

**Authors:** David V. Smith, James B. Wyngaarden, Sarah M. Weinstein, Nicholas Illenberger, Yinghua Liu, Jasmine Siegel, Bart Krekelberg

## Abstract

**Introduction:** Noninvasive brain stimulation can help clarify the neural basis of reward processing and potentially inform treatments for disorders involving reward dysfunction. However, widely used methods such as transcranial magnetic and electrical stimulation cannot directly stimulate deep-brain regions like the striatum. Here, we tested whether stimulating the ventrolateral prefrontal cortex (VLPFC)—a cortical region strongly connected to the striatum— could indirectly influence reward-related neural and physiological responses.

**Methods:** In a within-subjects design, participants performed a card-guessing task involving monetary rewards for correct guesses and punishments for incorrect guesses. During the task, participants underwent functional magnetic resonance imaging (fMRI) and pupillometry while receiving concurrent 10 Hz transcranial alternating current stimulation (α-tACS). Stimulation targeted either the VLPFC or a control region (temporoparietal junction). We measured pupil dilation, brain activation (BOLD signal), and functional connectivity between the ventral striatum and dorsal anterior cingulate cortex (VS–dACC).

**Results:** VLPFC stimulation increased pupil size during reward and punishment outcomes, indicating greater physiological arousal. At the neural level, α-tACS enhanced VLPFC activation during reward and suppressed its responses during punishment. Stimulation also changed VS– dACC connectivity in a context-dependent manner. Importantly, stimulation-driven increases in pupil size during reward correlated positively with stimulation-induced changes in VS–dACC connectivity. Exploratory moderated mediation analyses suggested that stimulation influenced the degree to which striatal responses mediated the relationship between task outcomes and pupil size changes.

**Conclusions:** Targeting VLPFC with α-tACS modulates local cortical activity and corticostriatal networks during reward processing, providing a promising noninvasive approach to influence reward circuitry.

**Highlights:**

- α-tACS over VLPFC increases pupil responses to reward and punishment.
- Stimulation alters reward-related VLPFC activity without enhancing striatal BOLD.
- α-tACS modulates ventral striatum–dACC connectivity in a task-dependent manner.
- Connectivity changes predict pupil dilation, linking brain and autonomic responses.

## 1. Introduction

Reward processing guides human decision-making, learning, and motivation. Central to this process is the striatal dopamine system, which encodes reward value, reinforcement signals, and adaptive behavioral responses [1, 2]. Functional neuroimaging studies consistently demonstrate that striatal activity scales with reward magnitude, integrating dopaminergic input to guide behavior [3, 4]. However, causal manipulation of reward-related neural responses in humans remains limited. Existing evidence predominantly comes from correlational fMRI studies, which reveal corticostriatal activation patterns but cannot confirm the necessity of these responses for behavior. This limitation is well-documented in neuroimaging critiques [5, 6], highlighting the need for causal manipulations of brain circuits to understand reward processing.

This gap is exacerbated by anatomical constraints, as standard noninvasive brain stimulation (NIBS) methods such as transcranial magnetic stimulation (TMS) and transcranial direct current stimulation (tDCS) cannot reliably target subcortical structures, such as the ventral striatum. Nonetheless, cortical stimulation may indirectly influence subcortical activity. For example, positron emission tomography (PET) studies have shown that tDCS over dorsolateral prefrontal cortex (DLPFC) induces dopamine release in both the ventral striatum [7] and prefrontal cortex [8]. However, these effects are variable, influenced by individual differences and baseline connectivity [9, 10]. Additionally, fMRI-tDCS studies indicate that prefrontal stimulation can modify perfusion in orbitofrontal cortex and caudate and alter connectivity between ACC and DLPFC during decision-making tasks [11]. Such variability underscores the need for precise, dynamic stimulation methods that modulate functional circuits in a state-dependent manner.

Transcranial alternating current stimulation (tACS) provides targeted modulation by entraining neural oscillations at task-relevant frequencies. Unlike constant-current tDCS, tACS delivers rhythmic sinusoidal stimulation, interacting with endogenous oscillations to influence large-scale networks selectively [12]. While most tACS research has examined cognitive control, perception, or resting-state connectivity, recent studies also demonstrate its potential in decision-making and learning contexts. For instance, frontal beta-frequency tACS increases risk-taking [13], while theta-frequency tACS modulates reversal learning performance [14, 15]. Still, the ability of tACS to influence reward-related corticostriatal dynamics during active task engagement remains unclear. Although tACS can alter cortical connectivity at rest [16], its impact on task-evoked corticostriatal interactions is largely unexplored.

In parallel, pupil dilation has emerged as a robust physiological marker of arousal and reward processing. Pupil responses reflect both dopaminergic and noradrenergic neuromodulation and correlate with reward magnitude, anticipation, and motivational salience [17, 18]. Recent evidence links blunted pupil responses to depressive symptoms [19] and shows that pupil dilation tracks subjective value and reinforcement learning [20]. However, whether neuromodulation alters pupil dilation during reward processing remains unknown. Notably, pupil dilation and alpha-band neural oscillations appear to reflect partly distinct neuromodulatory processes during cognitively demanding tasks [21], raising the possibility that these two signals could provide complementary insights into how reward circuitry is modulated.

Here, we applied alpha-frequency (10 Hz) tACS to the ventrolateral prefrontal cortex (VLPFC), a cortical region with strong anatomical projections to the striatum, to examine whether stimulation alters physiological, neural, and affective responses during reward consumption. We recruited 31 participants who completed a validated card-guessing task [22] during simultaneous fMRI, pupillometry, and tACS over VLPFC or an active control site (temporoparietal junction; TPJ), which falls outside the prefrontal-striatal circuitry targeted by our hypothesis [23]. After each block, participants rated their affective experience using the Positive and Negative Affect Schedule (PANAS). We addressed three primary questions: (1) Does α-tACS over VLPFC modulate affective, neural, and physiological responses to reward? (2) Does stimulation alter corticostriatal functional connectivity? (3) Are stimulation-induced neural changes linked to pupil responses during reward processing?

## 2. Methods

### 2.1. Participants

We collected neuroimaging data from 31 participants (20 males) aged 18 to 35 years (mean age: 23.45). The study was designed as an initial mechanistic test of α-tACS effects during concurrent fMRI and pupillometry; accordingly, the sample size was chosen to be feasible for this intensive within-subject protocol and consistent with prior concurrent stimulation–fMRI studies, rather than determined by a formal a priori power calculation. Participants were paid $25 per hour for their time. Participants were also paid for their performance on the reward processing task (see below). We excluded 3 participants for excessive head motion (details below), leaving a final sample of 28 participants (mean age: 23.35; 18 males) for all neuroimaging analyses. All participants provided written informed consent in accordance with the Institutional Review Board at Rutgers-Newark.

### 2.2. Reward Processing Task

Each participant completed 2–4 fMRI runs of a card guessing task that has been used extensively to study reward processing [22]. On each trial, participants viewed a mystery card and guessed whether its value was higher or lower than 5. Before the experiment, participants were informed that correct guesses were rewarded with $1.00, whereas incorrect guesses were punished with −$0.50. During the experiment, visual feedback informed them whether a guess was correct or incorrect. To maximize detection power, we used a blocked variant of the task similar to the Human Connectome Project [24]. Each block comprised eight trials (3.5 s per trial, including a 1-s intertrial interval; block duration = 28 s). The number of runs (2–4) was determined by session timing; participants completed as many runs as feasible within the scanning session. Concurrent with each block, we delivered 10 Hz transcranial alternating current stimulation (tACS) to either the ventrolateral prefrontal cortex (VLPFC) or a control site (temporoparietal junction; TPJ; see below). The stimulation site was assigned at the block level using a pseudorandomized, balanced schedule, so that participants received comparable numbers of VLPFC and TPJ blocks. Following each block of trials and stimulation, we assessed participants’ affective state using the Positive and Negative Affect Schedule [25] (see Supplementary Methods for details).

### 2.3. Transcranial Alternating Current Stimulation

We used a StarStim 8 device (Neuroelectrics, Barcelona, Spain) to generate tACS. We follow recommendations from Ekhtiari and colleagues [26] for safety and reporting procedures. Leads from the current generator in the control room connected to the shielding panel through radio-frequency filters (MRIRFIF; BioPac). Inside the scanner room, the custom, shielded cables contained a 5.6 k**Ω** resistor to limit RF heating and were wrapped in closed-cell polyethylene foam tubes to avoid overlapping wires and wire loops. Wires entered the bore from the side closest to the head and connected to the electrodes.

Subjects wore a neoprene headcap that allowed us to place the 8 stimulation electrodes (NG Pistim; 1 cm radius, Neuroelectrics, Barcelona, Spain). We used Signa electrode gel (Parker Laboratories Inc., Fairfield, NJ, USA) to ensure good connectivity between the electrode and the scalp, and tested impedances before entering the scanner, and again while lying in the scanner. All impedances were below 10 k**Ω**, most were below 5 k**Ω**.

The tACS waveform was a 10 Hz sinusoid with zero mean. Current amplitude ramped up and down linearly over a 2 s period at the beginning and end of stimulation. We used 3×1 montages to limit current spread. We targeted PFC by centering the montage over the expected location of the left VLPFC. The current amplitude at the center electrode (F3) was 750 **µ**A; current returned in equal proportions through three electrodes at FT7, Fp1, and FCz. In our active sham condition, we targeted the temporo-parietal junction (TPJ) with a montage centered on CP6 and current returns at CP2, PO8, and FT8. We selected TPJ as the active control site because our hypothesis specifically concerned prefrontal modulation of striatal reward processing, and TPJ is not a node in this prefrontal-striatal circuit [23]. This design also controlled for nonspecific effects of stimulation, including scalp sensation and arousal. The 3×1 electrode arrangement had a peak current density of 2.39 A/m^2^ at the center electrode. All participants tolerated the stimulation well; no sessions were aborted.

We used the ROAST package to simulate current flow based on these montages in the MNI-152 template brain (MNI152NLin6Asym). Following Huang et al., we set the conductivities as white matter: 0.126 S, gray matter: 0.276 S, CSF: 1.65 S, bone: 0.01 S, skin: -/456 S. All other parameters used the default settings defined in ROAST version 3.0.

### 2.4. Statistical Analysis of Behavioral Data

We used Matlab (R2021a, The Mathworks, USA), including its Statistics and Machine Learning Toolbox and several utility functions of our Matlab toolbox for linear mixed effects models (LMM) [27]. We specified each LMM using Wilkinson notation [28] and used maximum likelihood fitting procedures. All LMM included an intercept for each unique combination of subject and run as a random effect. For each term in the LMM, we report the F statistic, the degrees of freedom, and its associated p-value. We report effect sizes (and their 95% confidence intervals, CI) by expressing the relevant coefficient in the model as a percentage of the fixed-effect intercept. In other words, effect sizes for pupil size reflect percentage change relative to the grand mean pupil size, and effect sizes for PANAS scores reflect percentage change relative to the grand mean PANAS score.

### 2.5. Pupillometry

We tracked pupil size using an infrared eye tracker (Eyelink 2000, SR-Research, Canada) via a hot mirror aimed at the subject’s left eye through openings in the head coil. The eye tracker provided timestamps for blinks and a record of pupil size sampled at 500 Hz. For consistency, we analyzed only those trials and subjects that passed quality control and were included in the BOLD signal analysis (N=28 subjects). We determined the mean pupil size in the guess and outcome periods. In each period, we excluded the first 200 ms (to account for pupil latency) and a 300 ms interval around each blink (100 ms before and 200 ms after). Trials in which pupil tracking was not successful (e.g., due to excessive blinks or missing signal; 21% of trials) were removed from analysis.

During the guess period, participants waited for the appearance of the outcome card, which indicated whether they had guessed correctly. In reward blocks, the fraction of trials resulting in reward was higher (80%) than in punish blocks (20%). We hypothesized that this block structure and its associated higher expectation of reward could be reflected in the pupil size. To assess this, the LMM included a categorical predictor reflecting the block type (*rewardBlock*: true/false), and a categorical predictor to model the hypothesized effect of stimulation (*stim*: PFC/TPJ): *pupil∼ rewardBlock*stim + (1*|*subject:run)*.

At the start of the outcome period, the participant received visual feedback indicating whether they had guessed correctly. We hypothesized that this feedback could additionally modulate the pupil size (i.e., beyond the modulation based on the block structure). To assess this, we used a categorical variable (*correct*: true/false) and specified the LMM as: *pupil∼rewardBlock*correct*stim + (1 | subject:run)*.

### 2.6. Neuroimaging Data Collection

Neuroimaging data were collected at the Rutgers University Brain Imaging Center (RUBIC) using a 3 Tesla Siemens Trio scanner equipped with a 12-channel head coil. Functional images sensitive to blood-oxygenation-level-dependent (BOLD) contrast were acquired using a single-shot T2*-weighted echo-planar imaging sequence [repetition time (TR): 2.00 s; echo time (TE): 23 ms; matrix 68 x 68; voxel size: 3.00 x 3.00 x 2.50 mm; 36 axial slices (20% gap); flip angle: 76°]. To facilitate normalization of functional data, we collected high-resolution T1-weighted structural scans (TR: 1.9 s; TE: 2.52 ms; matrix 256 x 256; voxel size: 1.0 mm^3^; 176 slices; flip angle: 9°) and B_0_ field maps (TR: 402 ms; TE_1_: 5.19 ms; TE_2_: 7.65 ms; matrix 68 x 68; voxel size: 3.00 x 3.00 x 2.50 mm; 36 slices, with 20% gap; flip angle: 60°). In addition, we also collected T2-weighted structural images (TR: 3.2 s; TE: 494 ms; matrix 256 x 256; voxel size: 1.0 mm^3^; 176 slices; flip angle: 120°); these images are included with our data on OpenNeuro.org, but we did not use them in our preprocessing or analyses.

### 2.7. Neuroimaging Preprocessing and Analyses

Neuroimaging data were converted to the Brain Imaging Data Structure (BIDS) format using HeuDiConv [29] and minimally preprocessed using the default pipeline in fMRIPrep 20.1.0 [30], which builds upon Nipype 1.4.2 [31]. Full preprocessing details are provided in the Supplementary Methods. Following fMRIPrep, functional images were brain-extracted using FSL’s Brain Extraction Tool (BET) [32], spatially smoothed with a 6 mm full-width at half-maximum (FWHM) Gaussian kernel, and grand-mean intensity normalized using a single multiplicative scaling factor applied across the four-dimensional dataset.

All neuroimaging analyses were performed using FSL version 6.00 (FMRIB Software Library; www.fmrib.ox.ac.uk/fsl). Analyses were designed to evaluate whether ⍰-tACS modulated task-evoked brain activation and connectivity. Both activation and connectivity models were implemented within the general linear model (GLM) framework using FSL’s FEAT, with local autocorrelation correction enabled via FILM prewhitening.

To assess task-evoked activation, we specified a first-level model with four task regressors of interest: (1) reward trials with VLPFC stimulation, (2) reward trials with TPJ stimulation, (3) punishment trials with VLPFC stimulation, and (4) punishment trials with TPJ stimulation. Each regressor was convolved with FSL’s canonical double-gamma hemodynamic response function.

Task-dependent connectivity was assessed using generalized psychophysiological interaction (gPPI) models [33]. PPI is a widely used method that reveals consistent and specific connectivity patterns across psychological contexts and seed regions [34, 35]. We first identified seed regions based on two task contrasts: (1) ventral striatum, defined by the reward > punishment contrast, and (2) ventrolateral prefrontal cortex, defined by the interaction of stimulation site and task condition. For each participant, the mean time series within the seed region was extracted and entered as a physiological regressor, along with the four task regressors described above. Four interaction terms were generated by multiplying each task regressor with the physiological regressor, yielding 9 regressors (1 physiological, 4 task, 4 PPI) per gPPI model.

All first-level models included a common set of nuisance regressors: AROMA-derived motion components, the mean signal from white matter and cerebrospinal fluid, and indicators for non-steady-state volumes. High-pass filtering was implemented using a 128-second cutoff modeled with discrete cosine basis functions.

Functional data from multiple runs were combined within each participant using fixed-effects modeling. Group-level analyses were conducted using FMRIB’s Local Analysis of Mixed Effects (FLAME) stages 1 and 2 [36, 37]. Resulting z-statistic images were thresholded using a cluster-forming threshold of z > 3.1 and corrected for multiple comparisons using Gaussian Random Field Theory, with a cluster-level significance threshold of p < 0.05 [38, 39].

In exploratory analyses, we examined whether stimulation-related changes in brain activation and connectivity were associated with pupil responses. For each of the nine significant clusters identified in the group-level analyses, subject-level beta estimates were entered into robust linear regression models predicting pupil size, using the robustfit function in MATLAB. Each model included covariates for temporal signal-to-noise ratio (tSNR) and mean framewise displacement (fd_mean), both derived from MRIQC, to account for individual differences in data quality. To correct for multiple comparisons across clusters, p-values were Bonferroni-adjusted. We also conducted trial-level mediation analyses to test whether fluctuations in neural responses mediated the relationship between reward feedback and pupil dilation, and whether this mediation was moderated by stimulation condition. Full model specifications and implementation details are provided in the Supplemental Methods.

## 3. Results

### 3.1. Stimulation Alters Pupil and Affective Responses to Reward and Punishment

The pupil exhibited a characteristic response across each trial, with dilation beginning during the guess period and constricting following the outcome. VLPFC stimulation increased pupil size during both the guess and outcome periods, independent of reward context or task phase (see Supplementary Results and Figure S2).

Self-reported affect also varied across conditions, as assessed by the Positive and Negative Affect Schedule. Participants reported increased positive affect and reduced negative affect following reward blocks, consistent with the valence of recent outcomes. α-tACS over VLPFC had no significant effect on either positive or negative affect compared to TPJ stimulation (PANAS; see Supplementary Results and Figure S3).

### 3.2. Stimulation alters neural responses to reward and punishment

We began by testing our central hypothesis: that α-tACS over the ventrolateral prefrontal cortex (VLPFC) would modulate reward-related activation in the ventral striatum (VS). This hypothesis was grounded in the strong anatomical connectivity between VLPFC and VS and prior evidence that cortical stimulation can indirectly influence subcortical circuits. To test this, we conducted an ROI analysis focused on the reward > punishment contrast within bilateral VS (left VS: ke=136, x=-10, y=15, z=-1; right VS: ke=120, x=21, y=21, z=-4). A 2×2 repeated measures ANOVA revealed a significant main effect of valence, with reward outcomes eliciting significantly greater VS activation than punishment outcomes (*F*(1,25) = 33.13, *p* < .001, *η*^2^*p* = .57; Figure 2). However, there was no significant main effect of stimulation (*F*(1,25) = 0.04, *p* = .848, *η*^2^*p* = .001) and no significant valence × stimulation interaction (*F*(1,25) = 0.25, *p* = .618, *η*^2^*p* = .01). These findings suggest that α-tACS over VLPFC did not directly modulate striatal responses to reward during outcome processing.

**Figure 1.**
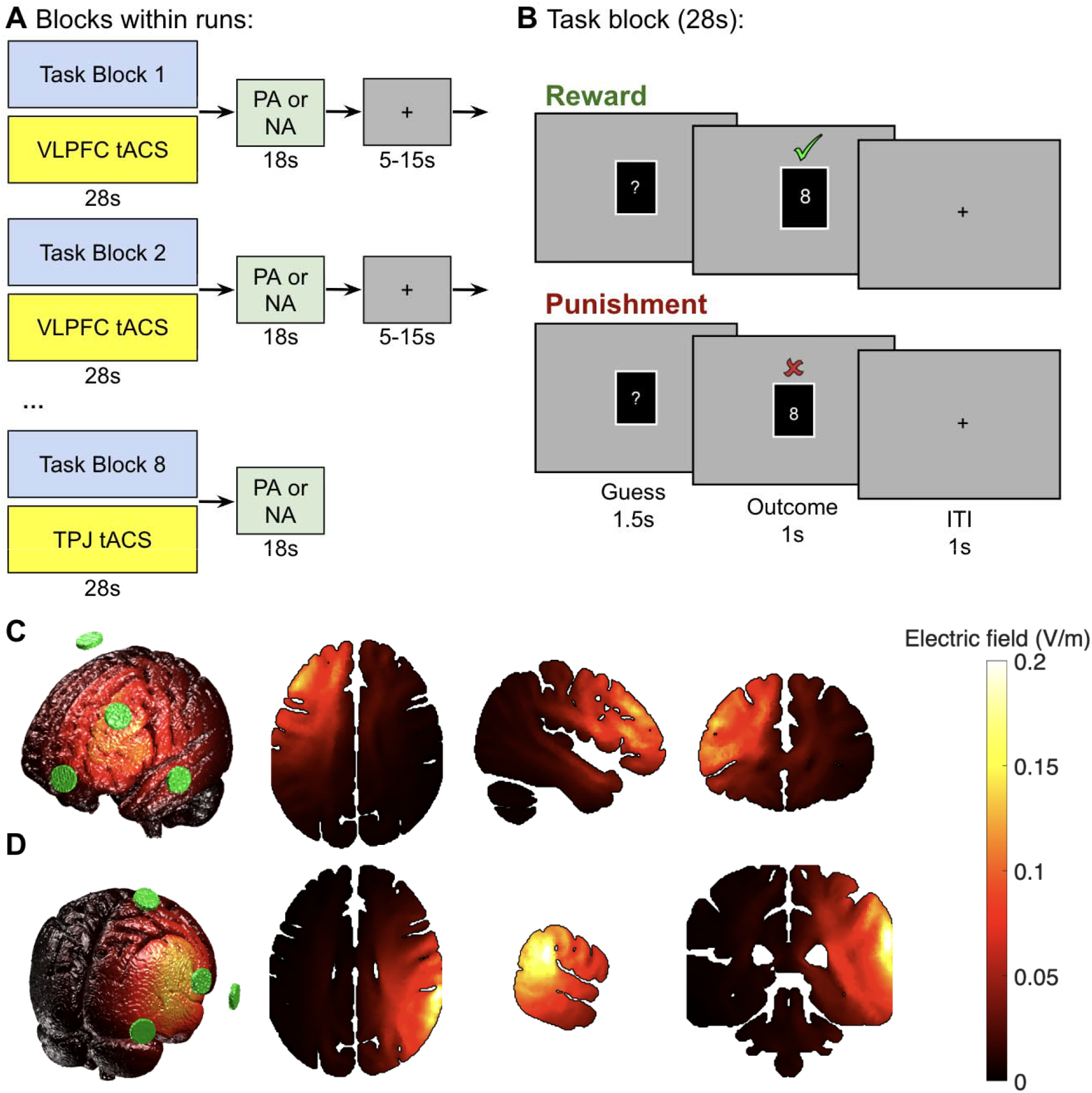
Experimental design and stimulation montages. (A) Run structure. Each run consisted of eight task blocks with concurrent α-tACS (10 Hz) targeting either the ventrolateral prefrontal cortex (VLPFC) or temporoparietal junction (TPJ; active control). Following each task/stimulation block (duration = 28s), participants completed Positive Affect (PA) or Negative Affect (NA) items from the PANAS (duration = 18s), followed by variable interblock fixation period ranging from 5 to 15s (mean=10s). Stimulation site varied across blocks within each run. (B) Trial structure of the card-guessing task. Participants guessed whether a hidden card value was higher or lower than 5, followed by feedback indicating correct (green checkmark) or incorrect (red X) responses. Each trial consisted of a guess period (1.5s), outcome period (1s), and intertrial interval (1s). Eight trials comprised each 28s task block that concurrently delivered stimulation. (C–D) Current flow simulations for the two stimulation conditions, generated using the ROAST toolbox. Green circles indicate electrode positions for the 3×1 montages. (C) VLPFC stimulation, with the center electrode at F3 and return electrodes at FT7, Fp1, and FCz. (D) TPJ stimulation, with the center electrode at CP6 and return electrodes at CP2, PO8, and FT8. Color scale reflects electric field magnitude (V/m).

**Figure 2.**
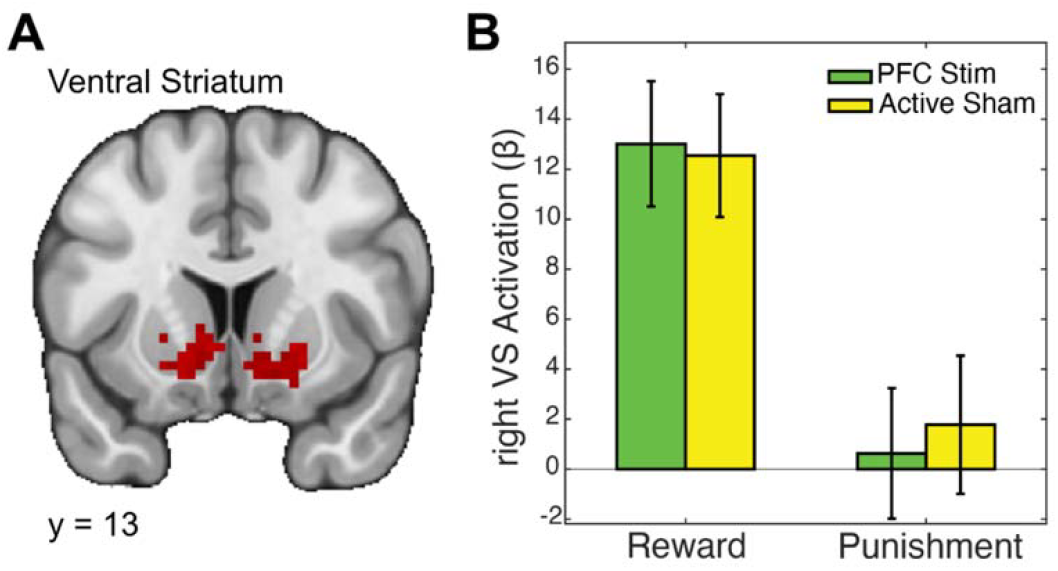
Ventral striatum responses to reward and punishment are unaffected by VLPFC vs. TPJ α-tACS. A) Whole-brain fMRI map highlighting bilateral ventral striatum (VS) activation clusters during the Card Guessing task. B) Bar graph showing beta estimates extracted from the right VS for Reward and Punishment conditions under 10 Hz transcranial alternating current stimulation (⍰-tACS) applied to the ventrolateral prefrontal cortex (VLPFC) or temporoparietal junction (TPJ; control, active sham site). Both stimulation conditions elicited robust VS responses to reward, with minimal activation during punishment. However, no significant differences were observed between VLPFC and TPJ stimulation in either condition, suggesting that VLPFC α-tACS does not directly enhance VS responsiveness to reward.

**Figure 3.**
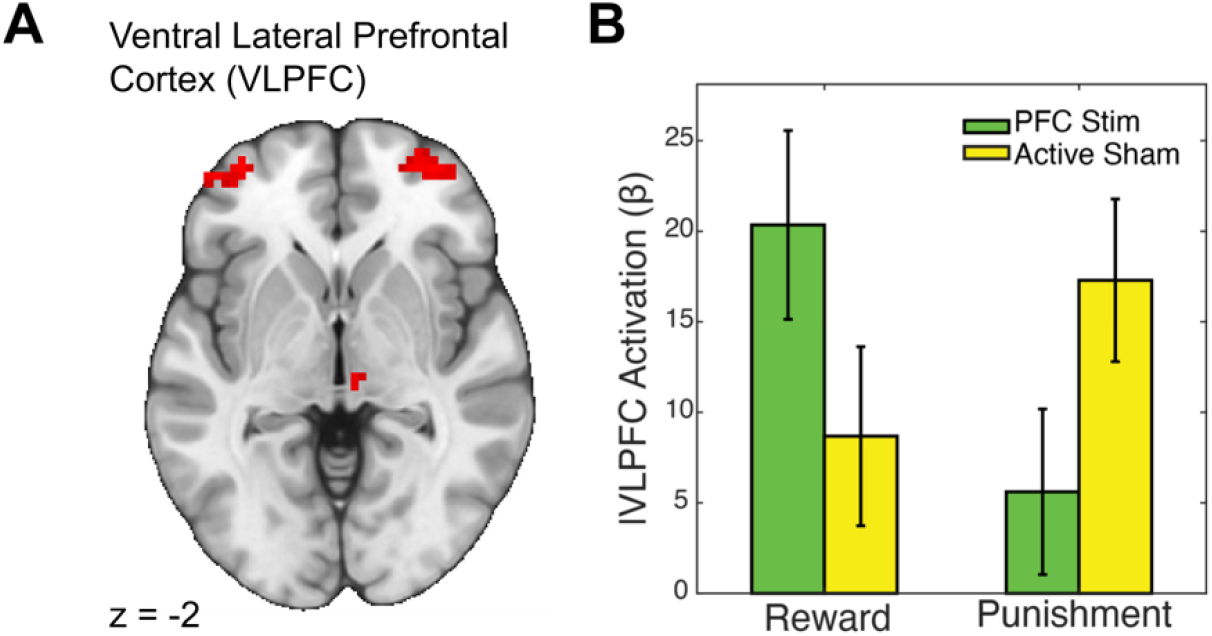
VLPFC α-tACS enhances local responses to reward and suppresses responses to punishment. Whole-brain fMRI map displaying bilateral ventrolateral prefrontal cortex (VLPFC) activation clusters during the Card Guessing task. B) Bar graph of beta estimates extracted from the VLPFC, comparing responses to Reward and Punishment conditions under 10 Hz α-tACS applied to VLPFC or temporoparietal junction (TPJ; control site). VLPFC stimulation significantly increased VLPFC activation during reward receipt compared to TPJ stimulation, whereas this pattern reversed during punishment, with TPJ stimulation eliciting greater VLPFC responses. These findings indicate that VLPFC α-tACS selectively modulates reward processing.

Although the predicted effects in the striatum were absent, a whole-brain analysis revealed compelling evidence for stimulation-induced modulation of cortical and subcortical responses. Specifically, we tested for interactions between stimulation site (VLPFC vs. TPJ) and task condition (reward vs. punishment) to identify regions where α-tACS produced outcome-specific effects. This analysis revealed significant clusters in bilateral VLPFC, as well as the thalamus, left inferior frontal gyrus, and supplementary motor area (see Table 1 for peak coordinates and cluster statistics). Visual inspection of beta estimates indicated that α-tACS over VLPFC increased local activation during reward and decreased activation during punishment, relative to the control stimulation site (Figure 4). These results provide converging evidence that VLPFC stimulation alters outcome-specific cortical responses, even in the absence of direct striatal modulation.

**Table 1.**
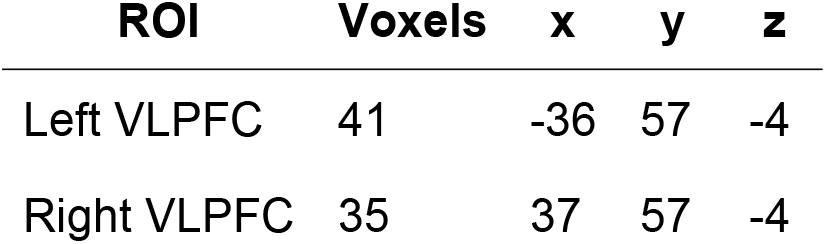
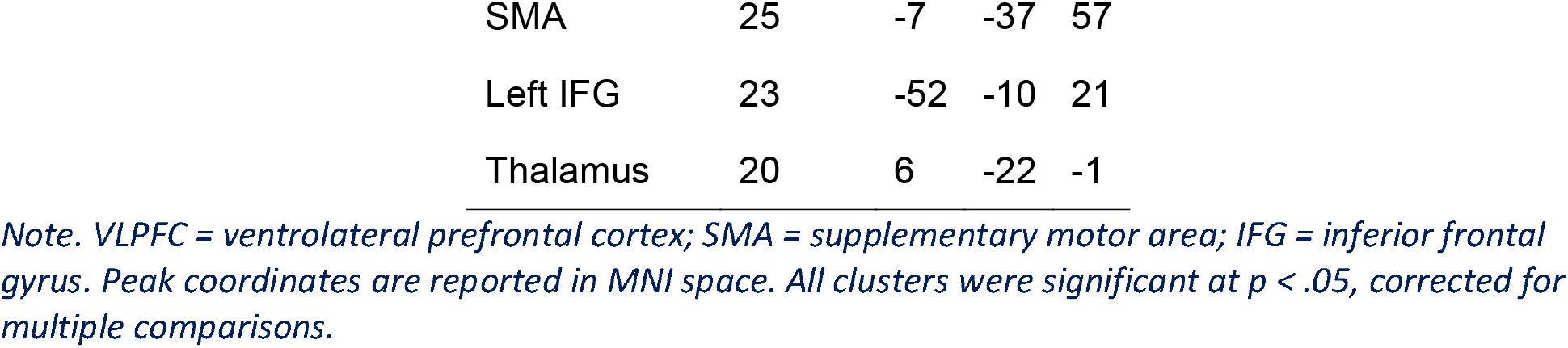
Activation Clusters for Reward > Punishment Contrast.

| ROI | Voxels | x | y | z |
| --- | --- | --- | --- | --- |
| Left VLPFC | 41 | -36 | 57 | -4 |
| Right VLPFC | 35 | 37 | 57 | -4 |
| SMA | 25 | -7 | -37 | 57 |
| Left IFG | 23 | -52 | -10 | 21 |
| Thalamus | 20 | 6 | -22 | -1 |

*Note. VLPFC = ventrolateral prefrontal cortex; SMA = supplementary motor area; IFG = inferior frontal gyrus. Peak coordinates are reported in MNI space. All clusters were significant at $p < .05$ , corrected for multiple comparisons.*

**Figure 4.**
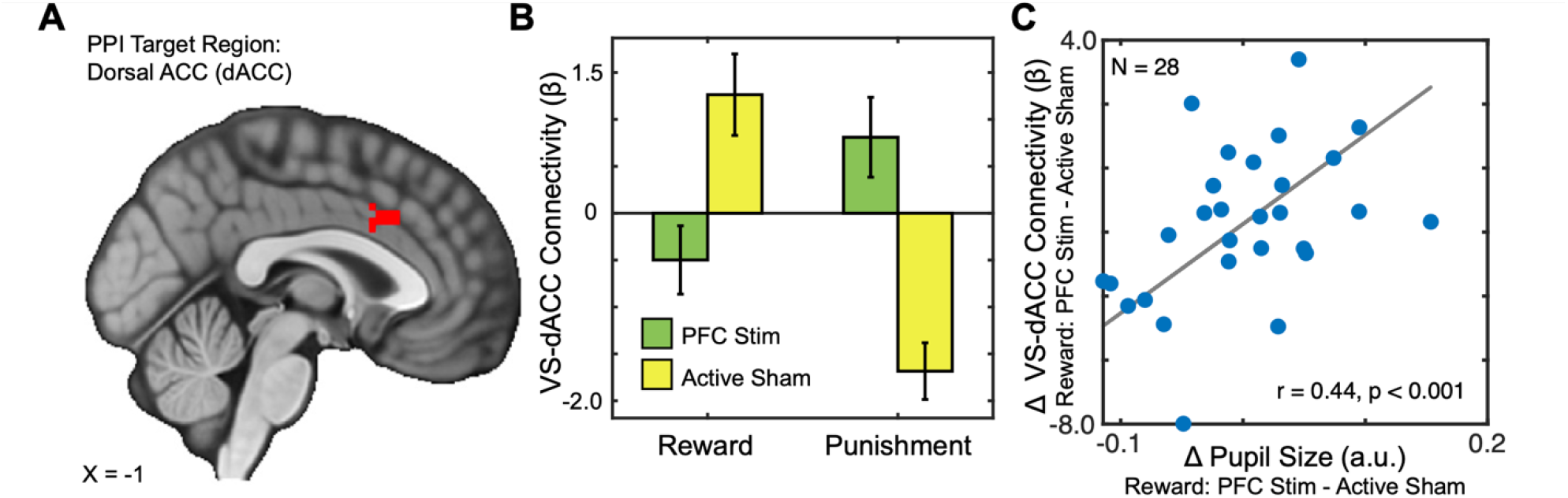
VLPFC α-tACS modulates ventral striatum-dorsal anterior cingulate cortex connectivity in a context-dependent manner. A) PPI target region in the dorsal anterior cingulate cortex (dACC). B) Bar graph depicting VS–dACC functional connectivity during Reward and Punishment conditions under 10 Hz α-tACS applied to the ventrolateral prefrontal cortex (PFC Stim) or active sham stimulation (TPJ; Active Sham). VLPFC stimulation dampened VS–dACC connectivity during reward receipt, contrasting with a pronounced positive increase under sham stimulation; this pattern reversed during punishment, suggesting that VLPFC ⍰-tACS modulates corticostriatal interactions in a context-specific manner, potentially disrupting reward-related network dynamics. C) Scatterplot showing a strong positive correlation (r = 0.44, p < 0.001) between reward-related changes in pupil size and VS–dACC connectivity, linking physiological and neural modulation. These results suggest that VLPFC α-tACS modulates corticostriatal networks in a context-specific manner, with potential implications for reward processing.

### 3.3. tACS Modulates Corticostriatal Connectivity and Pupil Responses

We next conducted functional connectivity analyses to determine whether α-tACS over the VLPFC altered task-evoked interactions between the ventral striatum (VS) and other brain regions. Given the central role of the VS in reward processing and its anatomical integration with cortical control systems, we conducted generalized psychophysiological interaction (gPPI) analyses using the right VS as a seed region. This analysis revealed a significant interaction between task condition (reward vs. punishment) and stimulation site (VLPFC vs. TPJ) on connectivity between the right VS and the dorsal anterior cingulate cortex (dACC; ke=220, x=3, y=21, z=36). Specifically, α-tACS over the VLPFC increased VS–dACC coupling during reward outcomes and reduced coupling during punishments, compared to TPJ stimulation. This effect was also observed in the precuneus (ke=350, x=-4, y=-61, z=54) and right angular gyrus (ke=190, x=12, y=-70, z=33).

To probe the functional significance of this connectivity effect, we next asked whether individual differences in VS–dACC coupling predicted pupil-linked arousal, an established physiological index of reward engagement. Indeed, across participants, the strength of VS– dACC connectivity was significantly associated with trial-averaged pupil dilation during reward outcomes (t = 3.887, p = 0.0007). This finding suggests that VLPFC stimulation modulates corticostriatal connectivity in a manner that tracks with enhanced physiological responsiveness to reward.

Finally, we sought to more directly test whether stimulation altered the pathway linking task condition to pupil responses via striatal activation. To do so, we implemented a high-dimensional moderated mediation analysis, assessing whether α-tACS moderated the relationship between task condition and local neural responses, which in turn predicted changes in pupil dilation. Strikingly, this analysis identified a distributed cluster of voxels within the striatum where task-evoked responses were modulated by stimulation in a manner that significantly mediated the effects of reward and punishment on pupil dynamics. These findings provide converging evidence that α-tACS influences arousal-linked neural pathways through specific modulation of corticostriatal circuits. (See Supplementary Methods and Results for full statistical details.)

## 4. Discussion

The present study used concurrent 10 Hz α-tACS, fMRI, and pupillometry to test whether rhythmic prefrontal stimulation can reshape reward-related circuit dynamics in humans. In a within-subject design with an active control site (TPJ), we found that VLPFC stimulation altered outcome-evoked cortical responses, shifted functional connectivity between the ventral striatum and distributed control systems, and modulated pupil-linked arousal. Because corticostriatal circuits support motivation, valuation, and adaptive behavior [1, 40], and because dysfunction in these systems is central to multiple psychiatric conditions [41-43], these findings help bridge a mechanistic gap: how noninvasive stimulation can causally influence reward-related brain dynamics during active engagement.

These results also clarify what α-tACS contributes relative to other neuromodulation techniques. TMS produces direct, suprathreshold cortical activation and is well-suited for causal perturbation, but its downstream effects on subcortical structures depend on circuit anatomy and current brain state. Transcranial ultrasound stimulation (TUS) offers greater depth and focality, enabling access to deep targets such as the striatum or nucleus accumbens, but its effects arise from mechanical rather than frequency-specific mechanisms [44, 45]. In contrast, tACS is optimized to interact with endogenous rhythms by biasing neural activity at specific frequencies, making it especially suited for modulating network coordination in a state-dependent manner. Prior work shows that alpha-frequency tACS can influence large-scale connectivity and affective outcomes, including anxiety, particularly when stimulation is aligned with an individual’s peak alpha frequency [46]. Recent clinical findings reinforce this idea, showing that stimulation outcomes vary with internal state and individual physiology, underscoring the potential for personalized protocols when targeting reward-related networks [47-49].

A key mechanistic insight from this study involves the dorsal anterior cingulate cortex (dACC), which is centrally involved in outcome monitoring, control allocation, and action updating. We found that α-tACS over VLPFC modulated connectivity between the ventral striatum and dACC in an outcome-dependent fashion, and that individual differences in this coupling tracked reward-evoked pupil dilation—a peripheral marker of arousal and dopaminergic tone. This finding aligns with prior work implicating striatal–cingulate interactions in reinforcement learning and behavioral adjustment [50-52]. It also connects to recent clinical research showing that frontostriatal connectivity, especially between the nucleus accumbens and anterior cingulate, predicts antidepressant response in major depression [53]. Rather than simply boosting striatal activation, α-tACS may modulate how ventral striatum and medial frontal regions coordinate during reward processing—an effect that is likely state-dependent and potentially sensitive to frequency alignment and individual targeting.

Despite these advances, several limitations merit discussion. First, the choice of 10 Hz alpha-frequency stimulation was theory-driven but did not test whether the results are frequency-specific. Prior evidence suggests that individualized stimulation frequencies — matched to intrinsic alpha or theta rhythms — may enhance neural and cognitive outcomes [49]. Future studies should systematically compare individualized and standardized protocols to better understand frequency-specific effects on reward circuitry. Second, reward processing encompasses distinct subprocesses, including anticipation, consumption, learning, and subjective valuation, which engage overlapping yet separable corticostriatal networks [54]. The present study specifically examined reward consumption; subsequent work should determine whether stimulation similarly modulates other reward subprocesses. Third, the absence of a sham or neutral control condition limits our ability to assess whether the observed effects reflect enhancement or suppression, as all contrasts are relative to active stimulation. This constraint weakens directional claims about causality, though our within-subject design and concurrent measurement of brain, pupil, and behavioral responses provide stronger inferential leverage than most stimulation studies [55].

It is also worth considering the sensitivity of different outcome measures to the stimulation parameters used here. Although α-tACS modulated both neural responses and pupil dilation during reward processing, we did not observe corresponding changes in self-reported affect. This dissociation is consistent with evidence that physiological measures are often more sensitive to neuromodulation than subjective report, and may reflect the relatively brief, single-session protocol employed. Relatedly, the short stimulation intervals (∼28 s per block) raise the possibility that longer or continuous protocols might yield more robust modulation of striatal activation. For clinical applications targeting affective dysfunction, these findings suggest that α-tACS can engage prefrontal-striatal reward circuitry at the neural level, but that higher doses or repeated sessions may be needed to produce changes in subjective experience. Future work should evaluate whether increased stimulation duration enhances neural modulation, autonomic sensitivity, and ultimately, affective and behavioral outcomes.

Collectively, these findings highlight α-tACS as a valuable method for investigating and modulating corticostriatal reward circuits. By demonstrating that brief, rhythmic prefrontal stimulation dynamically influences neural activation, functional connectivity, and autonomic markers of reward, this study adds critical mechanistic detail to our understanding of how oscillatory stimulation shapes reward processing. Concurrent neuroimaging and pupillometry provided converging evidence linking neural stimulation to physiological arousal, offering a promising methodological framework for future studies. Given that reward-processing abnormalities are central to numerous psychiatric disorders, including depression, bipolar disorder, and addiction, continued development of targeted neuromodulation strategies such as α-tACS could inform novel therapeutic approaches and deepen our understanding of transdiagnostic neural mechanisms underlying these disorders.

## Acknowledgments

This work was supported by the National Institutes of Health (R21-MH113917). DVS was a Research Fellow of the Public Policy Lab at Temple University during the preparation of this manuscript.

