## Supplementary Information for "α -tACS Modulates Reward-Dependent Pupil Responses and Corticostriatal Connectivity"

### 1. Supplementary Methods

#### 1.1. Block-to-block fluctuations in affect

To track changes in affective experience, subjects were prompted with three questions from the Positive and Negative Affect Scale (PANAS) [1] after each block. For instance: ‘At this moment, do you feel distressed?’. The time scale (‘at this moment’) was fixed throughout the experiment, but the word with affective connotations (e.g., ‘distressed’) was randomly chosen from the 20 words in the PANAS scale. Using the button box, subjects responded to the question with a response that varied in five steps from 1 (‘very slightly’) to 5 (‘extremely’). If a subject failed to respond within 6 seconds, the experiment moved on to the next question.

We summed the numeric responses in each block to obtain a positive (PA) and negative (NA) affect score and rescaled this to the original PANAS scale (which uses 10 probes). We hypothesized that PA and NA scores would be affected by the type of block immediately preceding the PANAS questionnaire and the stimulation applied during that block. We tested this hypothesis using a LMM with the following Wilkinson specification: *score~rewardBlock*stim+ (1|subject:run).* Here *rewardBlock* is a categorical variable with values true (indicating that the block preceding the PANAS question was a reward block) or false (a punish block). The *stim* variable is a categorical variable representing the stimulation target (VLPFC, or RTPJ). The *(1|subject:run)* term specifies a separate random effects intercept for each subject and run.

#### 1.2. Neuroimaging preprocessing

Neuroimaging data were converted to the Brain Imaging Data Structure [2] using HeuDiConv [3]. Results included in this manuscript come from preprocessing performed using *fMRIPrep* 20.1.0 [4], which is based on *Nipype* 1.4.2 [5]. The details described below are adapted from the fMRIprep preprocessing details; extraneous details were omitted for clarity.

##### 1.2.1. Anatomical data preprocessing

The T1w image was corrected for intensity non-uniformity (INU) with N4BiasFieldCorrection [6], distributed with ANTs 2.2.0 [7], and used as T1w-reference throughout the workflow. The T1w-reference was then skull-stripped with a *Nipype* implementation of the antsBrainExtraction.sh workflow (from ANTs), using OASIS30ANTs as target template. Brain tissue segmentation of cerebrospinal fluid (CSF), white-matter (WM) and gray-matter (GM) was performed on the brain-extracted T1w using fast via FSL 5.0.9 [8]. Volume-based spatial normalization to MNI152NLin2009cAsym standard space was performed through nonlinear registration with antsRegistration (ANTs 2.2.0), using brain-extracted versions of both T1w reference and the T1w template. To this end, the *ICBM 152 Nonlinear Asymmetrical template version 2009c* [9] template was selected for spatial normalization.

##### 1.2.2. Functional data preprocessing

For each of the BOLD runs, the following preprocessing steps were performed. First, a reference volume and its skull-stripped version were generated using a custom methodology of *fMRIPrep* [4]. Head-motion parameters with respect to the BOLD reference (transformation matrices, and six corresponding rotation and translation parameters) were estimated before any spatiotemporal filtering using mcflirt via FSL 5.0.9 [10]. BOLD runs were slice-time corrected using 3dTshift from AFNI 20160207 [11]. A B0-nonuniformity map (or *fieldmap*) was estimated based on a phase-difference map calculated with a dual-echo GRE (gradient-recall echo) sequence, processed with a custom workflow of *SDCFlows* inspired by the [epidewarp.fsl script](http://www.nmr.mgh.harvard.edu/~greve/fbirn/b0/epidewarp.fsl) and further improvements in HCP Pipelines [12]. The *fieldmap* was then co-registered to the target EPI (echo-planar imaging) reference run and converted to a displacements field map (amenable to registration tools such as ANTs) with FSL’s fugue and other *SDCflows* tools. Based on the estimated susceptibility distortion, a corrected EPI (echo-planar imaging) reference was calculated for a more accurate co-registration with the anatomical reference. The BOLD reference was then co-registered to the T1w reference using flirt via FSL 5.0.9 [10] with the boundary-based registration [13] cost-function. Co-registration was configured with nine degrees of freedom to account for distortions remaining in the BOLD reference. The BOLD time-series (including slice-timing correction when applied) were resampled onto their original, native space by applying a single, composite transform to correct for head-motion and susceptibility distortions. These resampled BOLD time-series will be referred to as *preprocessed BOLD in original space*, or just *preprocessed BOLD*. The BOLD time-series were resampled into standard space, generating a *preprocessed BOLD run in MNI152NLin2009cAsym space*.

Automatic identification of motion artifacts using independent component analysis (ICA-AROMA) [14] was performed on the *preprocessed BOLD on MNI space* time-series after removal of non-steady state volumes and spatial smoothing with an isotropic, Gaussian kernel of 6mm FWHM (full-width half-maximum). AROMA motion components were subsequently included as regressors in our analyses. Additional confounding time-series were calculated based on the *preprocessed BOLD*: framewise displacement (FD) and three regional signals (cerebral spinal fluid, white matter, and grey matter). FD was computed using the relative root mean square displacement between affines [10]. The three global signals were extracted within the CSF, the WM, and the whole-brain masks.

All resamplings were performed with *a single interpolation step* by composing all the pertinent transformations (i.e. head-motion transform matrices, susceptibility distortion correction when available, and co-registrations to anatomical and output spaces). Gridded (volumetric) resamplings were performed using antsApplyTransforms (ANTs), configured with Lanczos interpolation to minimize the smoothing effects of other kernels [15].

#### 1.3. Single-trial analyses

We estimated single-trial neural responses using the least squares single (LSS) approach, a variant of general linear modeling that isolates individual trial estimates while accounting for shared variance across trials [16]. For each trial, we constructed a separate GLM in which the trial of interest was modeled with its own regressor, while all other trials were grouped into a single nuisance regressor. Each model also included the same nuisance regressors used in our primary analyses, including motion parameters, framewise displacement, and physiological noise components. To account for serial autocorrelation in the residuals, all models incorporated local autocorrelation correction using FSL’s FILM prewhitening procedure. The resulting beta estimates provided trial-wise activation values for reward and punishment events under different stimulation conditions. These single-trial estimates served as inputs to our mediation analyses, enabling us to test whether stimulation-related changes in neural responses to feedback mediated changes in pupil dilation across trials.

#### 1.4. High-dimensional mediation analysis

We perform a two-step procedure to examine how (1) the effect of treatment (reward/punishment) on pupil size is mediated through brain activity, and (2) these mediation effects differ when brain stimulation targets either the RTPJ or VLPFC.

In the first stage of this procedure, partial least squares (PLS) is used to construct a set of features (i.e. linear combinations of voxels, or the columns of matrix M) that are maximally associated with the exposure (X1, reward/punish task), the moderator (X2, region of brain stimulation), and the interaction between the two, i.e.:

M ~ X1 + X2 + X1*X2	(Equation 1)

Where *M* is a matrix with dimension *r* x *p* of participant- and trial-level brain activity measurements (i.e., *r* is the sum of the total number of trials across all subjects and *p* is the number of voxels).

From the PLS model, we are able to derive a new set of features, $\tilde{M}_{k}=MV_{k}$, where $V_{k}$ is a *p*-dimensional vector whose elements map to brain voxels and thus forms linear combinations of voxels that maximize covariance between the left and right sides of Eq. (1). Below, when we interpret the average causal mediation effect (ACME) for the *k*th transformed mediator ($\tilde{M}_{k}$), we posit that this mediation is primarily driven by voxels corresponding to entries of $V_{k}$ with the largest weights.

These transformed features serve as the candidate mediators in our stage-two mediation analysis. In the second stage, we target the Average Causal Mediation Effect (ACME) of stimulation on pupil size working through each of the identified mediators. Specifically, using the potential outcomes framework [17], the ACME is defined as follows for mediators k=1,...,K:

$$\delta_{k}\left( 1 \right)=E[Y(1, \tilde{M}_{k}\left( 1 \right)-Y(1, \tilde{M}_{k}\left( 0 \right))]$$

For a specified mediator, $\tilde{M}_{k}$, we can conceptualize the amount of activity across voxels (weighted by $V_{k}$) if stimulation is targeted to the RTPJ ($M_{k}(1))$ versus if stimulation targeted to the VLPFC ($M_{k}(0)$). The ACME represents the average difference in pupil size that we would observe for a given participant if they received stimulation, but activity in this region were to be modulated between $M_{k}\left( 1 \right)$ and $M_{k}(0)$. In other words, the ACME represents the effect of stimulation that is specifically occurring due to changes in the highly weighted regions in $V_{k}$.

Mediation analyses were performed separately for each transformed mediator $\tilde{M}_{k}$ using the *mediate* package in R [18]. Because mediation effects were estimated one mediator at a time, residual dependence among the constructed mediators may bias ACME estimates; accordingly, we treat these mediation results as exploratory [19]. Causal mediation effects are nonparametrically identified under sequential ignorability assumptions (see [20]) and may be estimated given an input mediation model (region activity predicted by exposure) and outcome model (pupil size predicted by region activity and exposure). We assume that each mediator as well as the outcome can be modeled under the linear mixed effects framework with subject-level random intercepts. Specifically, mediators are modeled as a function of the treatment, moderator, and their interaction, and the outcome is modeled as a function of the candidate mediator, treatment, and a mediator-by-moderator interaction.

Estimated confidence intervals for mediation effects are limited in two important ways: (1) the mediation analysis does not take into account variability in the identification of mediators, and (2) because the mediators are identified using information from the full sample, observations may not be considered IID. In this situation, interpretation of confidence intervals must proceed under a “post-selection” framework. Specifically, conditional on a priori selection of the brain regions identified within stage one, inference for ACME in stage two is valid.

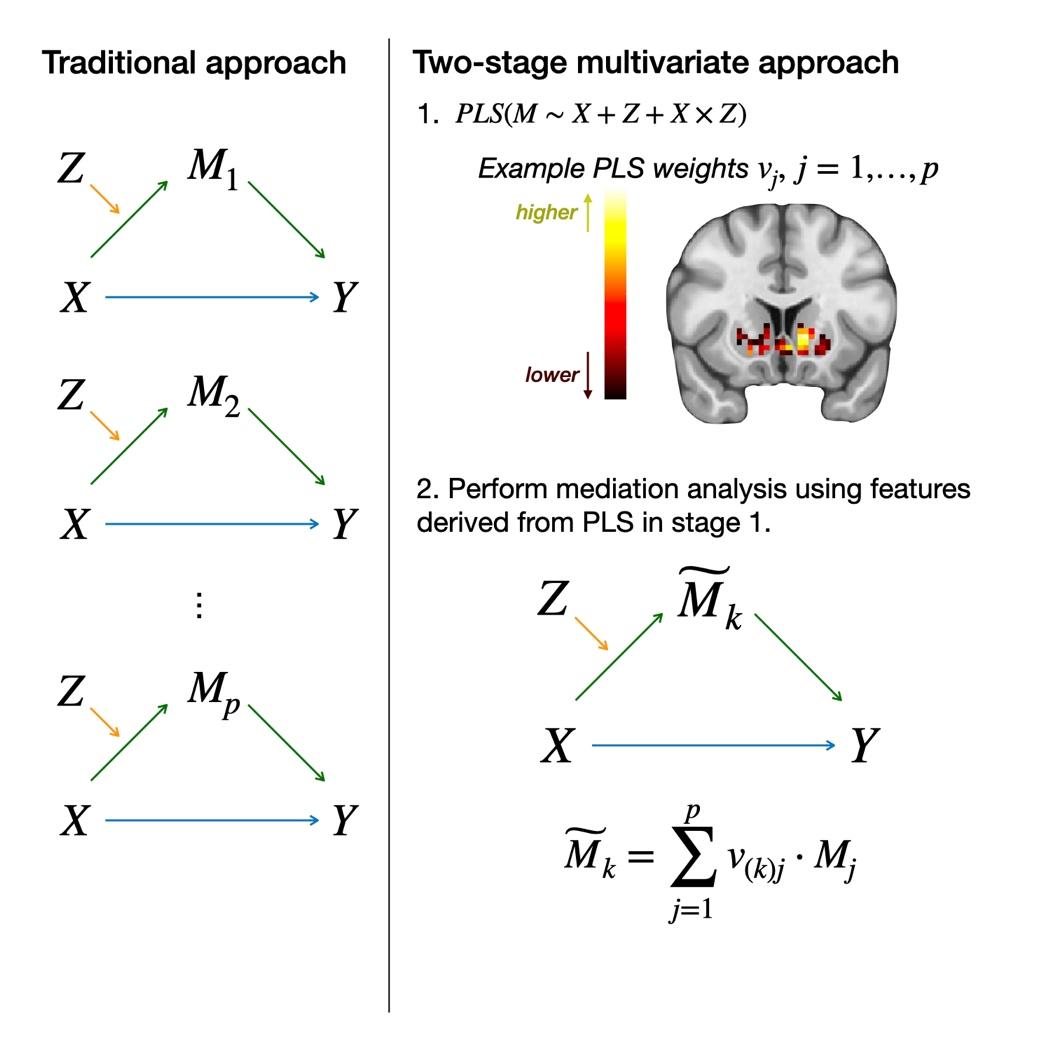

***Figure 1. Illustration of traditional vs. multivariate mediation approach.*** *Traditional mediation approaches in neuroimaging treat each voxel independently. In contrast, multivariate mediation approaches assess multiple voxels simultaneously. See text for analytic details.*

### 2. Supplementary Results

#### 2.1. Stimulation alters pupil responses to reward and punishment

The pupil exhibited a characteristic response across each trial (Figure 2). Dilation began during the guess period, peaking near the time of the participant’s response, and sharply constricted following the outcome before returning to baseline. During the guess period, pupil size was smaller in reward blocks (F(1,4654)=45.3, p=1.9e-11; Effect: -1.44%, CI [-1.86, -1.02])—consistent with reduced arousal when rewards were expected. VLPFC stimulation increased pupil size (F(1,4654)=16.8, p=4.15e-05; Effect: 0.88%, CI [0.46, 1.30]), and this effect was independent of reward probability (reward × stimulation interaction: F(1,4654)=0.03, p=0.85; Effect: -0.04%, CI [-0.46, 0.38]).

A similar pattern emerged during the outcome period. Pupil size was smaller in reward blocks (F(1,4562)=23.3, p=1.5e-6; Effect: -1.19%, CI [-1.67, -0.71])) and further reduced after correct guesses (F(1,4562)=257.1, p=2.5e-56; Effect: -3.96%, CI [-4.44, -3.47]). VLPFC stimulation again increased pupil size (F(1,4562)=8.51, p=0.0036; Effect: 0.73%, CI [0.24, 1.2]). No interactions among reward block, correctness, or stimulation reached significance (F(1,4562) < 2.1; p > 0.2). Together, these results indicate that pupil size reliably decreased under reward-predictive or rewarding conditions and increased with VLPFC stimulation, independent of reward context or task phase.

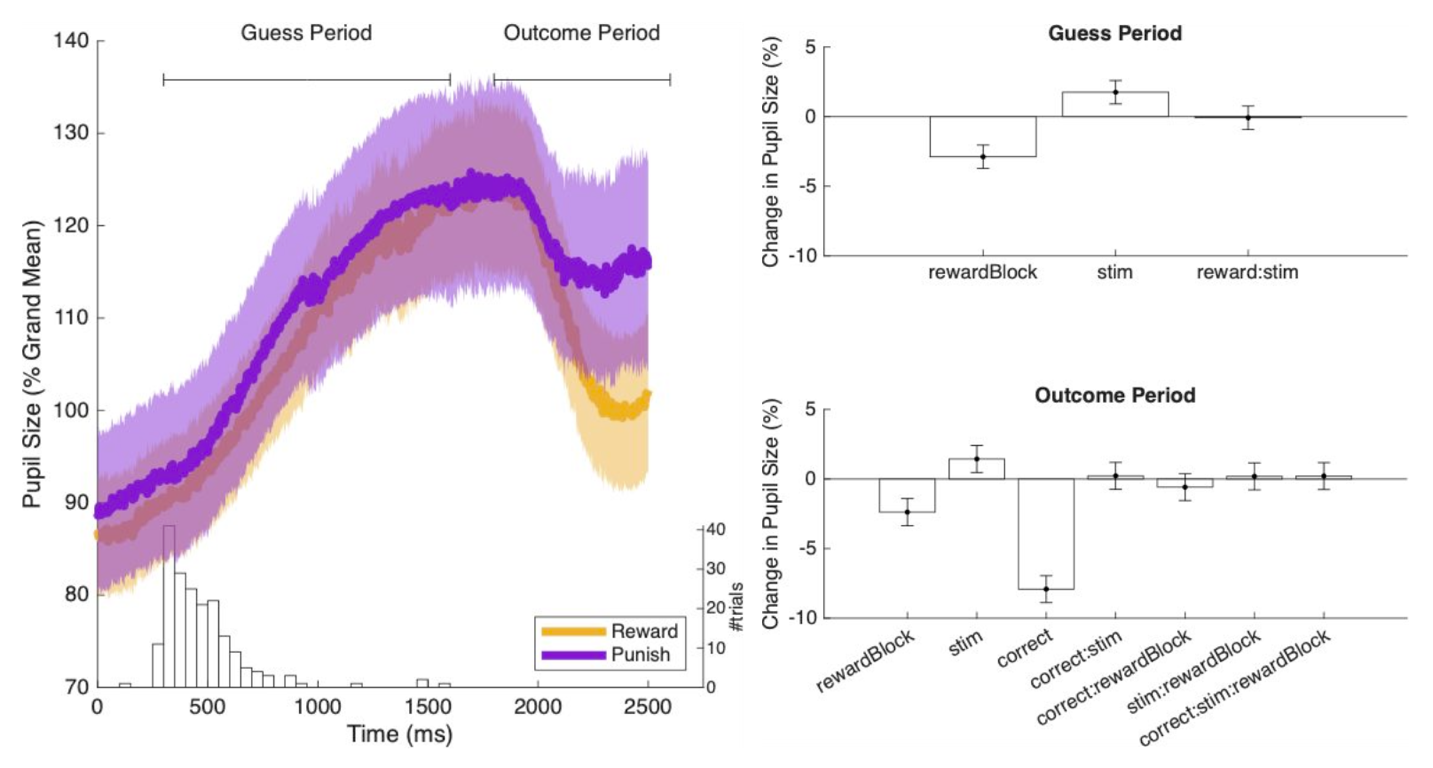

***Figure 2. Pupil size dynamics.*** *A) The curves show an example of the typical pupil size dynamics during the card game task (based on all trials of a single subject; ID#226). Traces were averaged separately for trials in reward blocks (yellow) and punish blocks (purple). Shading spans one standard deviation. The horizontal lines indicate the periods over which we averaged the pupil size to obtain a single measure of pupil size per trial, separately for the guess period and the outcome period. The histogram (with the y-axis on the right) shows the distribution of the times at which the subject pressed a button to indicate their guess. B) Analysis of the pupil size in the guess period. In this period, pupil size was significantly smaller in reward blocks and significantly larger when stimulation was targeted to VLPFC. C) Analysis of the outcome period. In the outcome period, pupil size was significantly smaller in the reward blocks and even smaller immediately after receiving a reward. VLPFC stimulation increased pupil size. Bars show the fixed effects for each of the model terms shown on the x-axis, error bars represent 95% confidence intervals. This figure shows that reward blocks and correct guesses elicited smaller pupil sizes, whereas VLPFC stimulation had the opposite effect.*

#### 2.2. Reward alters responses on the PANAS

For consistency, we included only PANAS questionnaire responses from blocks that were also included in the BOLD signal analyses. This yielded 2,528 responses from 28 subjects. Their overall positive affect (PA) score was 32.2 ± 9.9 (mean ± standard deviation), and their overall negative affect (NA) score was 18.7 ± 6.5 (Figure 1A). Both the PA score and the NA score were significantly larger than the normative scores of Watson et al (PA: 29.7 ± 7.9; p=0.0042; NA: 14.8 ± 5.4; p=1.1e-10, z-test). We suspect that this difference can largely be attributed to differences in the context where the PANAS scores were obtained: seated at a desk by Watson et al. or in the MRI scanner here.

Reward modulated PANAS measures of affective experience (Figure 3). PA was significantly higher after reward blocks (F(1,546)=25.2, p=7.14e-07, Effect: 4.52%, CI [2.75, 6.29]) and NA was significantly lower after reward blocks (F(1,554)=29.7, p=7.68e-08, Effect: -8.62%, CI [-11.7, -5.51]). Stimulation of the PFC had no significant effect on PA (F(1,546)=1.71, p=0.191, Effect: 1.18%, CI [-0.59, 2.94]), or NA (F(1,554)=0.00472, p=0.945, Effect: 0.109%, CI [-3.01, 3.23]).

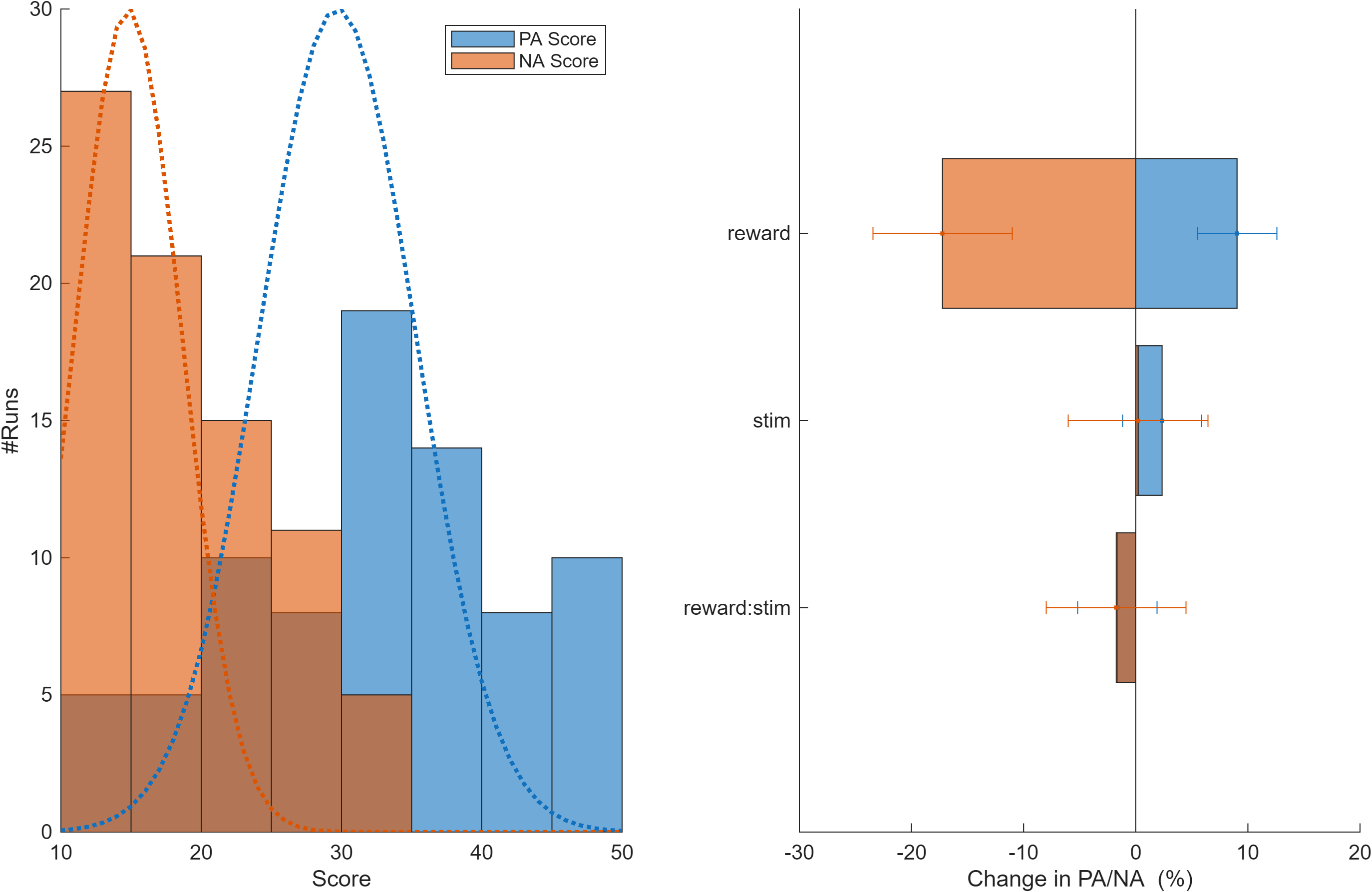

***Figure 3. Subjective affective experience as assessed by the Positive and Negative Affect Scale (PANAS).*** *A) Overall PA (positive affect) and NA (negative affect) scores based on 2528 probes across 28 participants. Dashed lines represent the large sample distribution observed by Watson et al. Our sample has higher PA and NA scores. B) Influence of reward, stimulation, and their interaction (reward:stim) as estimated by the LMM, expressed as a percentage of the grand mean PA and NA scores. Error bars show 95% confidence limits. Reward significantly reduced NA and significantly increased PA. α-tACS applied to PFC did not significantly modulate the PA or the NA.*

#### 2.3. α-tACS Moderates Striatal Mediation of Reward-Related Pupil Responses

To examine how alpha-frequency transcranial alternating current stimulation (α-tACS) influences the neural mechanisms linking reward feedback to autonomic arousal, we implemented a high-dimensional voxelwise moderated mediation analysis. Specifically, we tested whether α-tACS over the ventrolateral prefrontal cortex (VLPFC) moderated the indirect pathway from reward feedback (exposure, X) to pupil dilation (outcome, Y) through distributed patterns of brain activity (mediators, M_j_, indexed by voxel). This approach enabled us to evaluate whether stimulation alters the extent to which reward-evoked neural responses contribute to changes in autonomic function, operationalized via pupil size.

Results from this model identified clusters within the ventral striatum as the strongest contributors to moderated mediation effects (Figure 1). These findings suggest that α-tACS modulates how reward-related striatal activity translates into downstream autonomic responses. The spatial specificity of these effects—centered on canonical reward-related circuits—provides convergent evidence that α-tACS over VLPFC alters corticostriatal processing in ways that shape peripheral markers of motivational salience.
